# PYR1 biosensor-driven genome-wide CRISPR screens for improved monoterpenoid production in *Kluyveromyces marxianus*

**DOI:** 10.1101/2024.11.14.623641

**Authors:** Nicholas R. Robertson, Chase Lenert-Mondou, Alison C. Leonard, Aida Tafrishi, Stephanie Carrera, Sangcheon Lee, Yuna Aguilar, Leonardo Sanchez Zamora, Trinity Nguyen, Jesús Beltrán, Mengwan Li, Sean R. Cutler, Timothy A. Whitehead, Ian Wheeldon

**Affiliations:** Bioengineering, University of California, Riverside, Riverside, CA, USA; Biochemistry, University of California, Riverside, Riverside, CA, USA; Department of Chemical and Biological Engineering, University of Colorado Boulder, Boulder, CO, USA; Chemical and Environmental Engineering, University of California, Riverside, Riverside, CA, USA; Botany and Plant Science, University of California, Riverside, Riverside, CA, USA; Center for Industrial Biotechnology, University of California, Riverside, Riverside, CA, USA

## Abstract

Monoterpenoids are valued for their roles as flavors, fragrances, insecticides, and energy-dense fuels. Microbial biosynthesis offers sustainable biosynthesis routes for these important molecules, but production levels remain limited. Here, we introduce a biosensor-driven microbial engineering strategy to enhance monoterpenoid production, specifically targeting geraniol. Using mutagenized libraries of the PYR1 receptor—a versatile biosensor from plant ABA signaling pathways with a malleable binding pocket—we screened 24 monoterpenoids and identified PYR1 variants responsive to eight, including geraniol. A low background, highly selective geraniol-sensitive PYR1 variant was expressed in the thermotolerant yeast *Kluyveromyces marxianus* as a growth-based biosensor circuit, allowing for rapid strain engineering. By coupling the geraniol-sensitive PYR1 sensor with a genome-wide CRISPR-Cas9 mutagenesis approach, we identified six gene knockouts that enhance geraniol production, achieving up to a 2-fold increase in titer. This study demonstrates the power of the PYR1 biosensor platform to enable rapid strain engineering and the identification of mutants that improve titer of a desired metabolite.

## Introduction

Monoterpenoids are used as flavors, fragrances, insecticides, and high energy density fuels ^1–3^. Chemical production and plant extraction of these molecules often results in mixtures of terpenes that are costly to separate, rely on petroleum-based chemicals as inputs and in processing, and often result in low yields ^4–6^. An alternative to these processes is microbial biosynthesis which converts sugars or other renewable carbon sources into a desired product ^7^. Through synthetic biology approaches, microbial strains have been created that produce upward of a gram per liter of various monoterpenes ^8,9^; however, titers can be improved and other challenges such as production of unwanted byproducts by native enzymes still need to be addressed ^9,10^.

One successful approach to strain engineering is to couple a metabolite responsive biosensor to a quantifiable output, such as cell growth or the production of a fluorescent reporter ^11^. Such biosensor-driven screens can enable rapid identification of mutants that improve the production of a desired metabolite ^12^. Various biosensor technologies have been used for this, most often transcription factors ^13–16^, but sensors for many chemical classes are not available. There are a select few that have been developed for terpenes ^13,17^, but additional sensors are needed to enable rapid engineering of various industrially valuable monoterpenes.

PYR1 biosensors, from the plant ABA sensing system, are a flexible, sensitive, and specific platform ^18–20^. Ligand recognition occurs exclusively within the PYR1 receptor while the phosphatase partner, HAB1, acts as a co-receptor enabling high sensitivity and facile engineering of the binding pocket. This platform has been demonstrated in plants, bacteria, yeast, and ELISA-like *in-vitro* assays for classes of molecules such as natural and synthetic cannabinoids, agrochemicals, and environmental contaminants ^21,22^. PYR1 biosensors for terpenes have yet to be discovered and would be a valuable tool to improve the chemical scope of biosensor-driven screens.

To identify monoterpenoid biosensors, we screened three highly mutagenized libraries of PYR1 variants against a panel of 24 ligands and identified sensors for 8 monoterpenoids including geraniol, a flavor and fragrance molecule and the precursor to indole alkaloid pharmaceuticals and other high value metabolites ^23^. Porting these sensors into the fast-growing, thermotolerant yeast *Kluyveromyces marxianus* yielded functional GFP and growth-based reporting systems. We used the growth-based reporting system and a geraniol biosensor to build a biosensor-driven screening strain. Integration of geraniol synthase (*tCrGES*), the necessary heterologous step for geraniol biosynthesis, produced basal levels of geraniol to enable discovery of mutations that improve titers. We combined the PYR1 sensor strain with a genome-wide CRISPR screen to identify gene knockouts that improve geraniol production in *K. marxianus*.

## Results and Discussion

To isolate monoterpenoid activated PYR1 variants we screened three PYR1 mutant libraries against a panel of 24 different ligands (**Fig. 1**). Screening was accomplished using a yeast-two-hybrid (Y2H) approach ^24^; activation of PYR1 drives *URA3* expression enabling growth on selective media and isolation of sensor hits. Using this approach we screened three libraries of PYR1 variants. The first library was a computationally designed double-site mutagenesis (DSM) library with mutations focused in and around the PYR1 binding pocket. This library previously yielded a number of PYR1 variants receptive to a range of different ligands ^22^. A second library was created by shuffling first pass monoterpenoid hits from the DSM library with the original library (Terp-shuf.). We also computationally designed and constructed a new **h**ighly **m**utagenized library for hydrophobic ligands containing a single **H**-bond acceptor (HMH library). These libraries were screened for mutations that respond to monoterpenoids limited to three enzymatic steps from geranyl pyrophosphate (GPP): (+/-)-camphene, (+)-3-carene, (+)-fenchol, geraniol, (+)-(R)-limonene, (-)-(S)-limonene, (+/-)-(D)-linalool, (-)-ɑ-pinene, (-)-β-pinene, sabinene hydrate, ɑ-terpineol, which are one enzymatic step from GPP; carveol, citral, (+)-β-citronellol, D-(+)-fenchone, L-(-)-fenchone, (+)-limonene 1,2-epoxide, myrtenol, (-)-perillyl alcohol, ɑ-pinene oxide, and (S)-cis-Verbenol, which are two enzymatic steps from GPP; and, camphor, (R)-(-)-carvone, and (S)-(+)-carvone, which are three enzymatic steps from GPP.

**Fig. 1.**
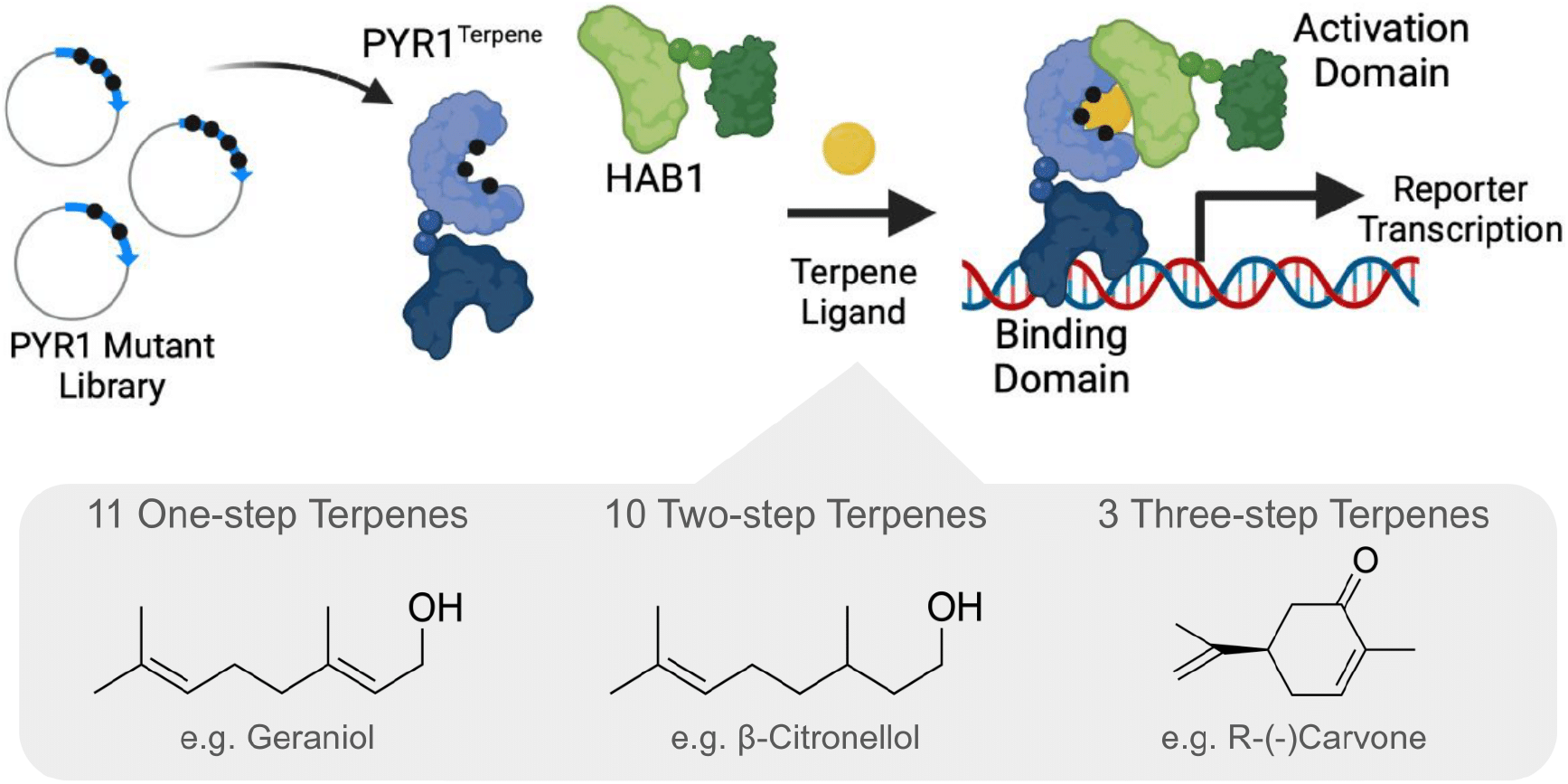
Metabolic pathway and screening for terpene biosensors. Terpenes relevant to metabolic engineering were screened for active PYR1 receptors. Mutant PYR1 receptor libraries were screened to find mutated binding pockets that accept terpenes. PYR1 is bound to a DNA binding domain and the dimer partner of PYR1, HAB1, is bound to a DNA activation domain. When the appropriate ligand is present, PYR1 and HAB1 reconstitute activating transcription of a reporter gene. We screened well-studied terpenes one, two, or three enzymatic steps away from the natively produced precursor, GPP. In total we screened 11 one-step, 10 two-step terpenes, and 3 three-step terpenes.

In total, we identified 50 distinct receptor sequences for 8 monoterpenes ((R)-(-)-carvone, β-citronellol, geraniol, (+/-)-linalool, myrtenol, (-)-perillyl alcohol, ɑ-terpineol, and (S)-cis-verbenol). Seven of the hits were from the DSM library, 10 from Terp-shuffle, and the remaining 33 from HMH (**Fig. S1)**. Binding pocket mutations of the most sensitive hits to each ligand and the chemical similarities of the ligands are presented in **Fig. 2a**. Sensitivity data and the chemical structure of each hit is shown in **Fig. 2b**. When used as biosensors to drive a genetic circuit, these monoterpene responsive PYR1 mutants can be used in biosensor-driven strain engineering. Here, we investigated a set of 15 sensors responsive to geraniol to create strains of the yeast *K. marxianus* to overproduce geraniol.

**Figure 2.**
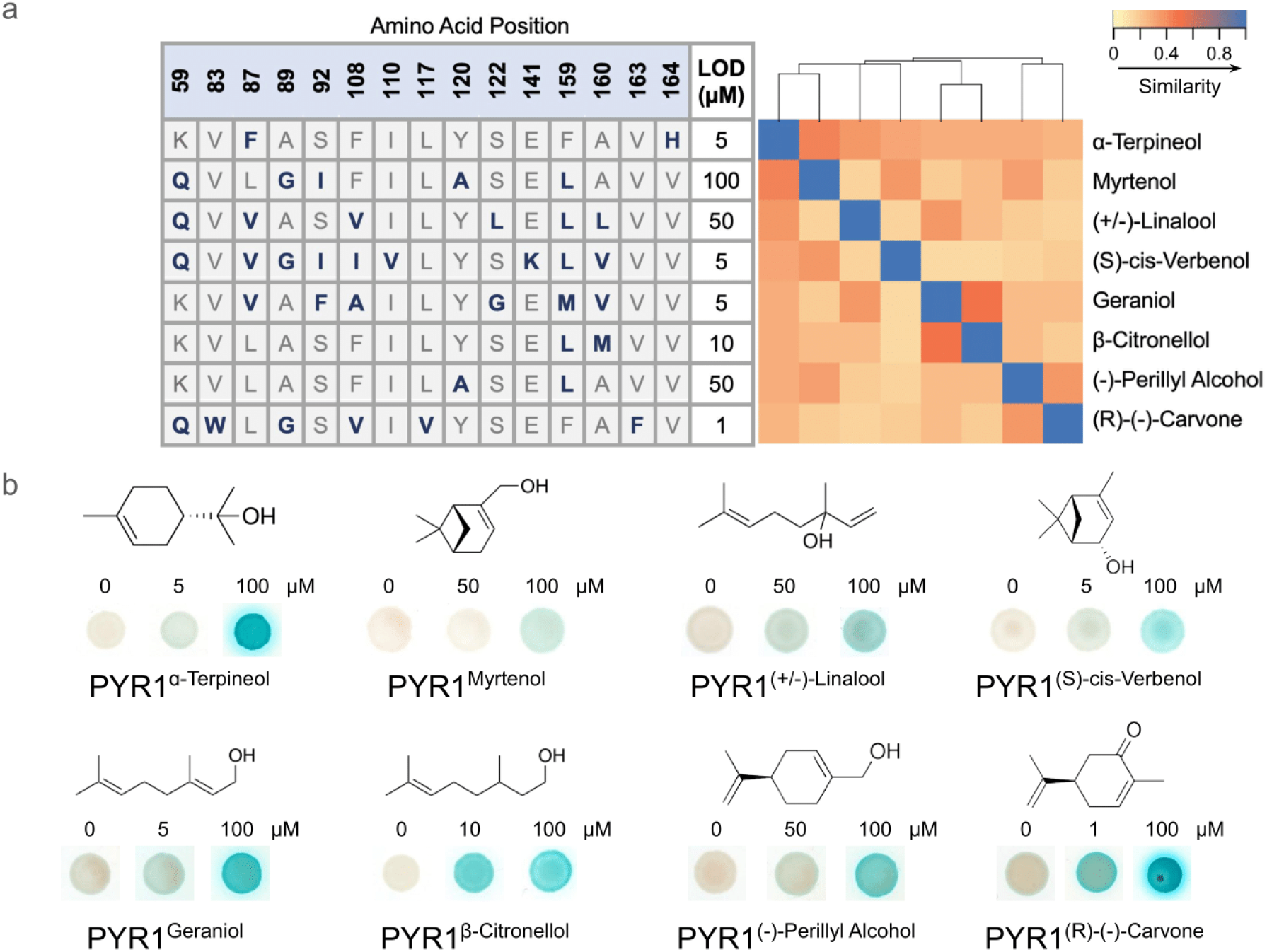
Sequences and sensitivity of PYR1 mutants sensitive to monoterpenoids. (a) Amino acid substitutions for our lowest sensitivity receptors for each ligand. Blue letters indicate an amino acid substitution. Black numbers in the top row indicate the amino acid position in the PYR1 binding pocket. The heat map indicates chemical similarity based on chemical clustering using ChemMineR ^25^. (b) Chemical structure of monoterpenes with Y2H blue/white X-gal staining of *S. cerevisiae* containing the respective PYR1 mutant. Blue indicates activation of the receptor by the on-target monoterpene. DMSO control (0 μM), the lowest visible blue staining (or 50 μM if the sensitivity is 100 μM), and 100 μM concentration of each ligand is displayed.

Geraniol was chosen as the target monoterpene because it is a precursor to many valuable compounds, but also because we isolated a number of PYR1 variants with different properties for this compound, thus providing a number of options from which we could build the strain engineering pipeline. Six geraniol-responsive receptors with varying sensitivity were ported to *K. marxianus* as genetic circuits and tested for sensitivity, selectivity, background signal, and fold-activation. Cross-reactivity against geranic acid and β-citronellol was tested due to their structural similarity and because they are natural degradation products of geraniol (**Fig. S2**) ^9^. Of the six characterized receptors, one was selected as the best for screening because it has low background response, low cross-reactivity with the geraniol degradation products, and μM sensitivity as shown in **Fig. 3a**. We call this PYR1 variant, PYR1^geraniol^.

**Figure 3.**
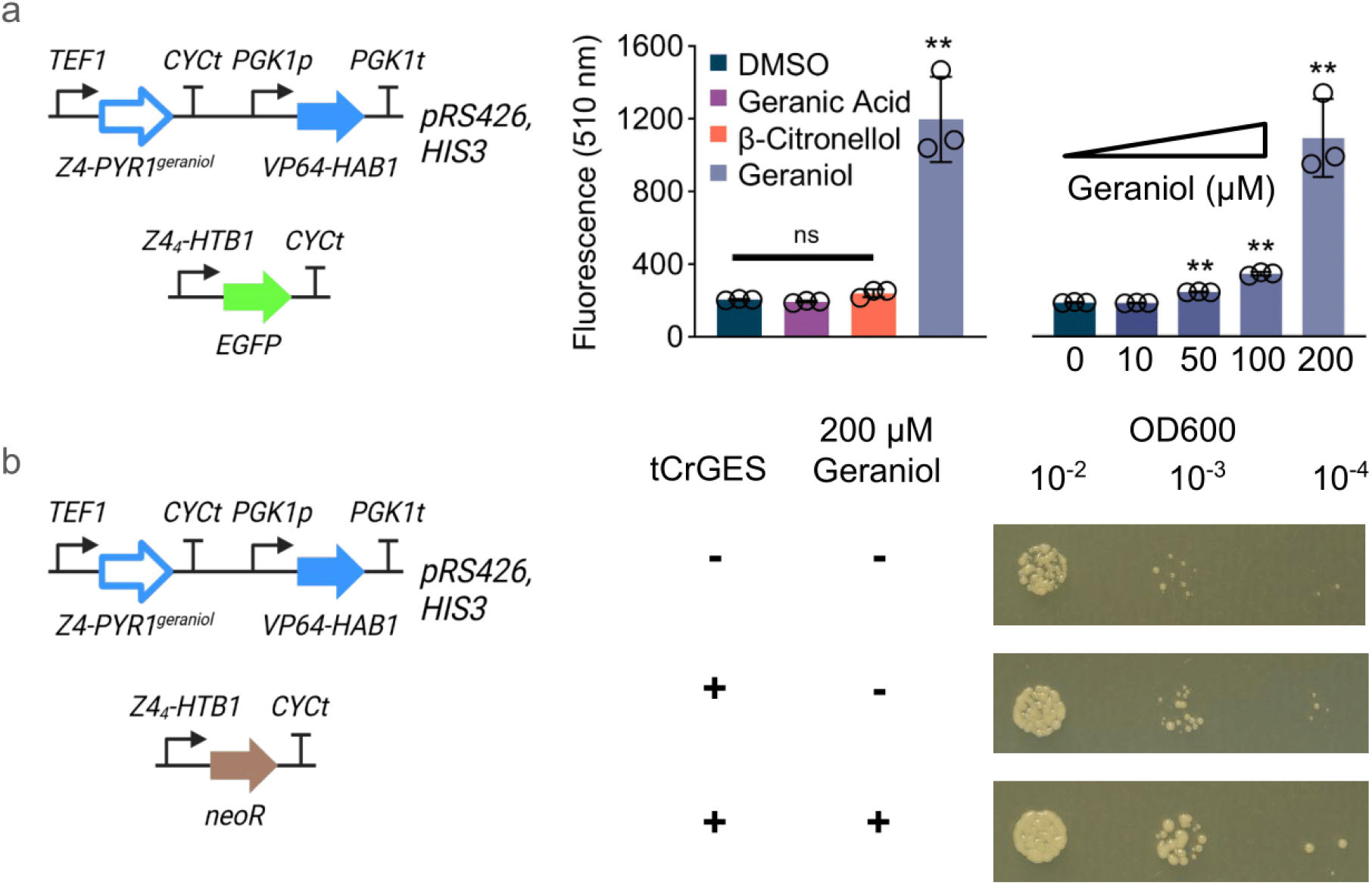
PYR1 monoterpene biosensors function as reporters in *K. marxianus*. GFP fluorescence and cell growth on G418 plates as reporters for biosensor activation in *K. marxianus*. (a) Gene circuit and fluorescence data for activation of PYR1^Geraniol^. The PYR1/HAB1 cassette is carried on a replicating plasmid and the GFP reporter is integrated at the *ABZ1* locus. Sensor response was tested with 10, 50, 100, and 200 μM geraniol added to the media, while geranic acid and β-citronellol were tested at 200 μM with DMSO as a negative control. There was no significant (ns) cross reactivity with the degradation products and the sensor responded to 50 μM geraniol (** indicates p<0.005; n = 3). Overnight cultures were used to inoculate 1 mL of SD-H liquid culture to an OD600 of 0.05. After 12 hours at 30 °C and 1,000 RPM shaking, fluorescence was measured at 510 nm by flow cytometry. (b) Growth spot assays of *K. marxianus* harboring a plasmid expressed PYR1^Geraniol^ genetic circuit driving G418 resistance (*NeoR*). Cells were grown overnight in SD-H liquid culture then spotted on solid SD-H media containing 3 mg/mL G418 and incubated at 30 °C for 30 hrs. *K. marxianus* CBS6556 with *URA3Δ::Z4*_*4*_*-NeoR I4::CAS9*, and PYR1^Geraniol^/HAB1 contained on a plasmid with and without XYL2::*tCrGES* were characterized.

Next, we tested whether responsiveness to geraniol can be used in a selection assay to enable screening for mutants that improve geraniol biosynthesis. We sought to test geraniol-dependent growth on media containing the antifungal agent G418 with the PYR1^geraniol^ circuit driving *neoR* expression which provides resistance to G418 (**Fig. 3b**). The *URA3* marker used in *S. cerevisiae* was not suitable here as the sensor expression plasmid for *K. marxianus* already utilizes this marker. Growth on the selection condition improved over the baseline when geraniol synthase (*tCrGES*) was integrated into the *K. marxianus* genome; this heterologous gene is required for geraniol biosynthesis in *K. marxianus*. To simulate the screening condition and improve geraniol biosynthesis, we added 200 μM geraniol to the media, which produced more growth than base levels of geraniol production via the integration of *tCrGES* alone and significantly more growth than the control strain. Given the properties of this biosensor and the geraniol-dependent growth observed we chose this sensor and strain for our subsequent screens.

With our growth-based screening system in place, we can now screen a mutant library for variants that improve geraniol production. To generate the mutant *K. marxianus* library, we used a genome-wide CRISPR sgRNA library with 55,614 unique guides. This library targets every annotated protein coding region (CDS) in the genome with 10-fold coverage (*i*.*e*., 10 unique guides per gene). The final strain is a stable haploid (*Δalpha3 Δkat1) K. marxianus* CBS6556 with *HIS3Δ URA3Δ::Z4*_*4*_*-NeoR XYL2::tCrGES I4::CAS9*, and PYR1^Geraniol^/HAB1 expressed from a replicating plasmid. With Cas9 expressed from a genome integrated expression cassette and selection based on antifungal resistance, this strain enabled growth-based genome-wide screening. Transformation of our sgRNA library into our screening strain yields a pool of mutants, which contain single knockouts that improve geraniol production (**Fig. 4a**). The screen was performed on solid media because geraniol is known to be rapidly exported from the cell potentially leading to a high number of cheater cells in a liquid based screen (*i*.*e*., cells respond to geraniol in the media and not geraniol produced intracellularly ^9,11^).

**Figure 4.**
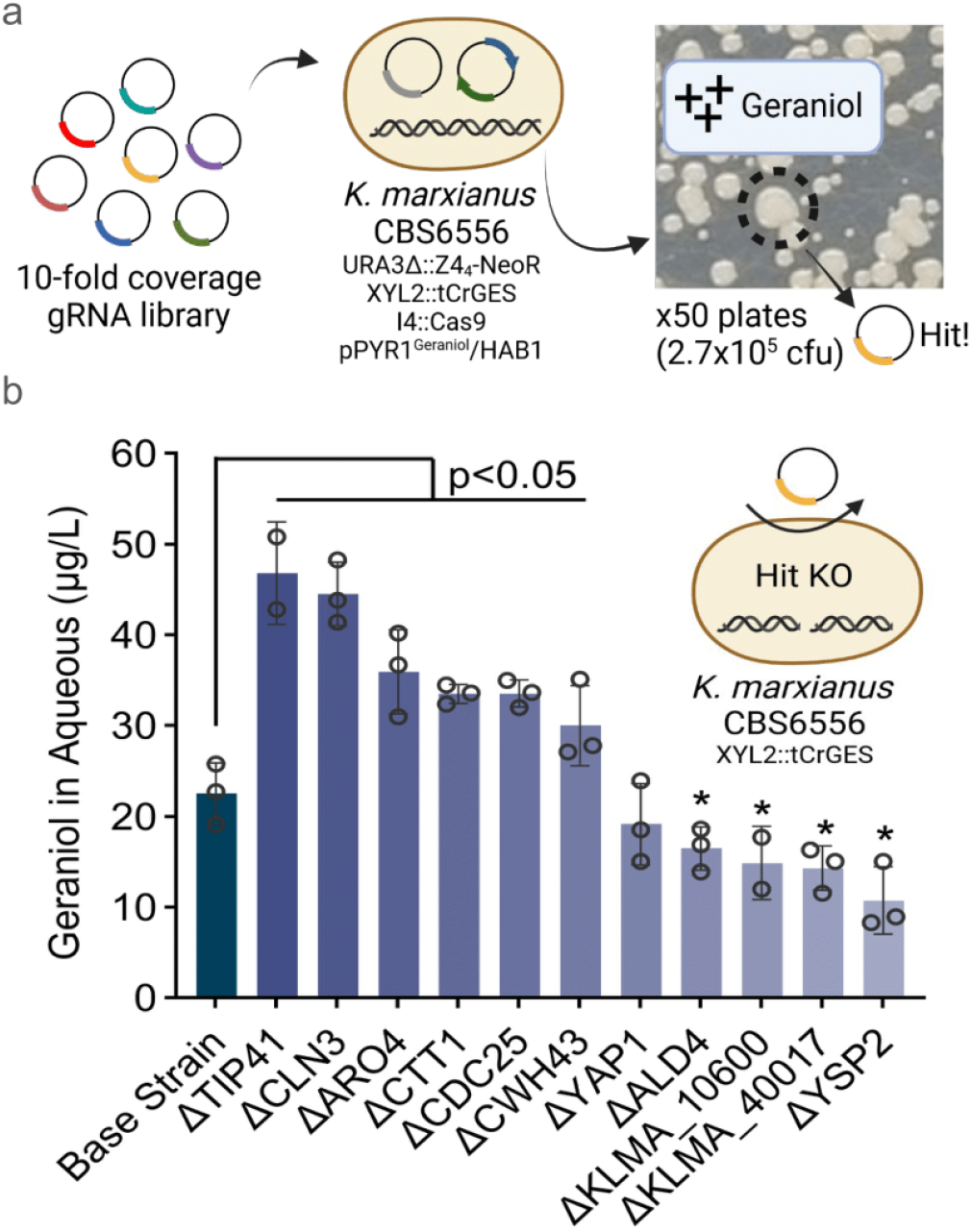
Biosensor-driven genome-wide screens reveal knockouts that improve geraniol production. Hits from the growth-based screen were investigated and evaluated for terpene production. (a) A 10-fold coverage genome-wide library was transformed into our growth-based screening strain with HAB1/PYR1^Geraniol^ on a plasmid. If a knockout improves terpene production, strong growth will be observed on selective plates. (b) Geraniol production of gene knockout hits. Six hits exhibited significantly (p<0.05) more geraniol production than the base strain, while four hits (* represents p<0.05) produced less geraniol. Samples were taken after 16 hrs of growth in 250 mL shake flasks with 50 mL YPD and 2 mL dodecane overlay grown at 220 RPM, 30 °C. The dodecane layer was extracted and ran on GC-FID to extrapolate geraniol production in the aqueous phase.

In total, 2.7×10^5^ cells were plated on selective media, representing nearly 5-fold coverage of the sgRNA library. Colonies produced from these cells were visually inspected to identify the largest colonies, which theoretically represent mutations that improved geraniol biosynthesis. We observed two important results from the screen: (1) that roughly half of the plated cells survived on G418 plates; and (2) that ∼1% of the colonies were visibly larger than others (**Fig. 4a**). We selected a subset of the larger colonies for further analysis and validation. Of those tested, we identified 11 strains that contained a CRISPR-induced edit to the target gene that introduced a premature stop codon, thus disabling gene function (**Fig. S3**). To validate these hits, gene knockouts were remade in a strain containing only the integrated *tCrGES* cassette. Six of the hits improved geraniol production, two of which improved production by two-fold (**Fig. 4b**). Four others decreased geraniol production and one putative hit had no effect on geraniol titer. In total, we observed a majority (∼55%) successful hit rate. This diversity of screen results is consistent with our previous work ^26–28^ and from other biosensor-driven strain engineering pipelines ^29,30^.

Two of the validated hits provide a clear link between gene function and geraniol biosynthesis. For example, *CTT1* is known to play a key role in terpene tolerance in *S. cerevisiae* ^*31*^ and when knocked out has been shown to increase β-caryophyllene in *S. cerevisiae* ^32^. *ARO4* is a critical component of the Shikamate pathway which is highly active in *K. marxianus*, which produces key precursors to 2-phenylethanol and a corresponding ester ^33,34^. These products divert carbon away from geraniol biosynthesis and are toxic to the cell in high quantities ^34^. Knockout of *ARO4* reduces carbon flux to these products and may also reduce overall oxidative stress allowing for additional geraniol production. The four remaining gene knockouts that improve geraniol titers are *TIP41* (negative regulator of the TOR signaling pathway in *S. cerevisiae*) ^35^, *CLN3* (involved in vacuolar arginine transport and pH homeostasis in *S. cerevisiae*) ^36^, *CDC25* (A phosphatase with activity in cell cycle progression in *S. pombe*) ^37^, and *CWH43* (A cell wall maintenance protein in *S. cerevisiae*) ^38^. These hits represent new strain engineering targets for improving monoterpenoid biosynthesis in yeast.

PYR1 biosensors are a flexible, portable, and specific platform for synthetic biology. We screened highly mutagenized PYR1 libraries for monoterpenoid biosensors and found mutants that are activated by 8 different ligands, including geraniol, a precursor to many valuable metabolites. When fused with DNA binding and transcriptional activation domains PYR1 and HAB1 can be used to create genetic circuits for controlling gene expression, thus enabling genetic screens for improved production of geraniol. With this goal in mind, we ported a low background, high fold-activation PYR1^geraniol^ circuit to the fast growing yeast *K. marxianus*. To create genetic diversity in *K. marxianus*, we combined a sensor expressing strain with a genome-wide CRISPR sgRNA library and selected for mutants that improved growth in a growth-based selection. The screen identified a series of gene knockouts that improved geraniol production, two of which improved geraniol titer by 2-fold over the base strain. Overall, this work shows that coupling novel PYR1 sensor discovery with genome-wide mutagenesis enables high-throughput strain engineering for a target metabolite.

## Materials and Methods

### Microbial Strains and Culturing

All strains, plasmids, primers, and sgRNAs used in this work are listed in **Tables S1, S2, S3, and S4**, respectively. Notably, *S. cerevisiae* strain BY4742 *ura3Δ his3Δ LeuΔ TrpΔ* was utilized as the base *S. cerevisiae* strain, while *K. marxianus* strain (CBS6556 *Δalpha3 Δkat1 ura3Δ his3Δ)* was used as the base *K. marxianus* strain. Strains without plasmids were grown in YPD media (10 g L^−1^ yeast extract, 5 g L^−1^ peptone; DB Difco®, Becton-Dickinson, 20 g L^−1^ glucose). All yeast strains harboring plasmids with auxotrophic markers were cultured with synthetic defined (SD) media minus the selective amino acid. For example, strains containing plasmids with a uracil auxotrophic marker were cultured in SD-U media: 6.7 g L^−1^ BD Difco™ Yeast Nitrogen Base without amino acids (Sigma-Aldrich), 1.92 g L^−1^ Yeast Synthetic Drop-out Medium Supplements without uracil (Sunrise Science Products), and 20 g L^−1^ D-glucose. *K. marxianus* and *S. cerevisiae* strains were grown at 30 °C. Liquid cell cultures in shake flasks were grown at 300 rpm in a shaker incubator. Liquid cell cultures in 96 deep-well plate formats were grown at 990 rpm in a shaker incubator.

### Yeast Two-Hybrid Screening of Mutant Biosensor Libraries

Selection experiments for mutant receptors that respond to new ligands were conducted as previously described by Beltrán^22^. Briefly, in this Yeast two-hybrid (Y2H) system, PYR1 is fused to the GAL4-DNA binding domain (BD) and HAB1 is fused to GAL4-DNA activation domain (AD). Upon ligand binding, PYR1 and HAB1 form a stable complex bringing together the BD and the AD. This activates transcription from promoters harboring a GAL4 responsive UAS. MAV99 (MATa trp1-901 leu2-3,112 hisΔ200 ade2-101 gal4Δ gal80Δ can1rcyh2rLYS2::(GAL1::HIS3) GAL1::lacZ SPO13::10xGAL4site::URA3) was used as it allows for direct selection for and against PYR1 mutants that enable ligand-induced HAB1 interactions by GAL4-mediated transcription of its URA3 cassette. Expression of the URA3 marker allows for both positive selections (via rescue of uracil auxotrophy) or negative selections, by sensitivity to the uracil anti-metabolite 5-fluoroorotic acid (5-FOA). The pBD-PYR1 mutant libraries were transformed into MAV99 harboring pAD-HAB1. After transformations, negative selections were conducted to remove receptors that bind HAB1 in a ligand-independent fashion (i.e., constitutive receptors) by growing the library on solid media containing 1 g/L 5-FOA; the purged library was collected and used in subsequent selections for cells responsive to 200 μM terpenoid ligands on SD-Trp,-Leu,-Ura media. Colonies supporting uracil-independent growth at 30 °C were isolated after 3 days, regrown on SD-Trp,-Leu,+1 g/L 5-FOA to screen out constitutive biosensors, retested to confirm ligand-dependent growth on SD-Trp,-Leu,-Ura plates with and without test chemical, and finally validated by β-galactosidase staining and growth validation.

We used three mutagenized PYR1 libraries to screen for sensors: a previously described computationally designed double site-saturation mutagenesis (DSM) library^22^, a new library constructed by using DNA shuffling to diversify four initial terpene sensors isolated from the DSM library with the DSM library (Terp-shuf.), and a new, highly mutagenized, computationally designed library (HMH). Details on the library computational design and construction by Golden Gate assembly^39^ can be found in the Supplementary Information file and in **Tables S5-S6**. For each library, ≥1 million colony forming units (cfu) were screened per library per terpene.

### Genome-Wide Knockout Screening

Sixteen library transformation reactions were pooled and inoculated into 500 mL of SD-HU media. Samples were collected during mid log phase (24 hours, OD_600_ = 5.4) and early stationary phase (28 hours, OD_600_ = 7.6). Glycerol stocks were prepared from culture taken at these time points with a final concentration of 30% glycerol and stored at -80 °C. OD600 was also measured at 44 hrs and found to be 8.0.

To identify knockouts that are beneficial to terpene production, glycerol stocks of cells were plated on SD-HU + 3 g/L G418 at a concentration of 1,000-10,000 cfu/plate. After incubation at 30 °C for 40 hrs, a large diversity of colony size was observed. Approximately ½ of plated colonies survive on G418 plates and of these about 1% are obviously much larger. The largest colonies were picked and streaked on SD-HU+G418. An individual colony from this streak was used to identify the gRNA contained within the cell. If the guide was active, amplification and sequencing of the targeted gene was performed to find if a knockout occurred.

## Data availability

SRA deposition # to be added at publication. All 8 of the most sensitive biosensor plasmids for each monoterpenoid (**Fig. 2**) and PYR1^Geraniol^ will be made available from Addgene.

## Author contributions

NRR, SRC, and IW conceived the project. NRR, CLM, and JB conducted the biosensor screen and characterized isolated mutants. ACL and TAW designed and built the highly mutagenized PYR1 library. AT and ML designed and constructed the sgRNA library. NRR, SC, YA, SL, TN and LSZ optimized library transformations, conducted the CRISPR screens, and validated hits. NRR, CLM, and IW wrote the manuscript. All authors edited the manuscript.

## Acknowledgments

This work was supported by DARPA CERES, NSF-2225878, NSF-2128016, NSF-2128287 (T.A.W.), and an NSF GRFP Award #1650115 to A.C.L.

