## Supplemental Materials for "PYR1 biosensor-driven genome-wide CRISPR screens for improved monoterpenoid production in *Kluyveromyces marxianus*"

Table S1. Strains
Table S2. Plasmids
Table S3. Primers
Table S4. Guide RNAs
Table S5. HMH Library Design
Table S6. HMH Library Coverage
Figure S1. All Monoterpenoid Biosensors
Figure S2. GFP Reporting of 6 Geraniol Biosensors in *K. marxianus*

### Materials and Methods (Extended)

### Yeast Transformations

For *K. marxianus,* a single colony was picked and grown in 2 mL YPD for 24 hrs at 30 °C. This starter culture was used to inoculate 50 mL of YPD in a 250 mL shake flask at a starting OD600 of 0.01. After growing to an OD600 of 15.5-16.5 (~16 hrs at 30 °C), 2 mL of cells were harvested by centrifugation at 4000 g for 1 minute. Cells were washed once with 0.1 M lithium acetate, 1X TE. To the cell pellet, 10 µL of ssDNA (Agilent, 300-2000 bp, 10 µg/µL) was added followed by 800 ng plasmid DNA (800 ng of each plasmid for dual plasmid transformations) and then 500 uL transformation buffer (39% PEG4000, 0.15 M lithium acetate, 30 mM dithiothreitol). This solution was mixed well by pipetting and allowed to rest at room temperature for 15 minutes. Heat shock was performed for 7 minutes at 47 °C. One milliliter of water was added and the transformation was centrifuged at 4000 g for 1 minute. After disposing of the supernatant, cells were resuspended in 1 mL of sterile water. The transformed cells were then plated on selective plates to calculate transformation efficiency and/or grown in liquid culture to allow for gene knockouts or integrations. Our single plasmid transformations into Ys626 yield up to $5x10^5$ transformants per reaction.

For *S. cerevisiae*, a single colony was picked and grown in 2 mL SD-Leu overnight at 30 °C. Next, 1 mL of this starter culture was used to inoculate 50 mL of SD-Leu in a 50 mL conical tube. After growing to an OD600 of 1.0-1.2 (14-16 hrs at 30 °C), the cells were harvested by centrifugation at 2,000 g for 5 minutes. Cells were resuspended in 20 mL 5X TE, LiOAc (0.5 M, pH 7.5) and shaken at 200 RPM, 30 °C for 45 minutes. Then, 500 uL of dithiothreitol (DTT) was added to the solution and incubated for an additional 15 minutes. Next, 30 mL ice-cold sterile $H_2O$ was added and the solution was centrifuged at 2,000 g, 4 ℃ for 5 minutes. The supernatant was removed and the cell pellet was resuspended in 50 mL cold $H_2O$, the mixture was then centrifuged at 2,000 g, 4 ℃ for 5 minutes. The supernatant was again removed and the pellet was resuspended in 1 mL of cold $H_2O$. This was then transferred to a pre-chilled 1.5 mL tube and centrifuged at 6,000 g, 4 ℃ for 1 minute. The supernatant was removed and 1 mL ice-cold 1 M sorbitol was added, mixed, and centrifuged at 6,000 g for 1 minute at 4 ℃. The supernatant was again removed and the cell pellet was resuspended in 400 µL ice-cold 1 M sorbitol. These cells were used at competent cells and were stored at 4 ℃ for up to a week. To transform, 80 µL of these competent cells were combined with 1.5 µg of plasmid DNA and mixed well. This mixture was then incubated on ice for 15 minutes. The mixture was then transferred to a 0.4 cm ice-cold electroporation cuvette and pulsed at 2.5 kV, 25 µF, 200 Ω. Immediately 1 mL ice-cold 1 M sorbitol was added and the mixture was incubated for 30 minutes at room temperature. Finally the mixture was grown on SD-Trp,-Leu containing 1 M sorbitol for 3 days.

### Molecular Cloning and Reagents

All primers used in this work are listed in Table S3. All generated gene construct sequences and plasmid maps are provided as Multimedia Components 1-38. Cloning reagents and restriction enzymes were purchased from New England Biolabs (NEB). All primers, guides, and gBlock DNA fragments for DNA cloning were purchased from Integrated DNA Technologies (IDT). All guide sequences used in this work are listed in Table S4. The Q5® High-Fidelity DNA polymerase system was used for DNA amplification. NEBuilder® HiFi DNA Assembly Master Mix was used for Gibson assembly. All PCR products and linearized vectors were fractionated by agarose gel electrophoresis and purified using a Zymo Research gel extraction kit. All plasmids were propagated in *E. coli* TOP10 cells (Thermo Fisher Scientific), and plasmid extractions were performed with the Zymo Research plasmid miniprep kit. All g-block DNA fragments used in this work are provided in Table S5.

### Monoterpenoids

Biosensor strains were induced with the following chemicals: (+/-)-Camphene (Cayman Chemical, CAT#: 24478); Camphor (Sigma-Aldrich, CAT#: 148075-100G); (+)-3-Carene (Cayman Chemical, CAT#: 23165); Carveol (Cayman Chemical, CAT#: 33831); (R)-(-)-Carvone (Fisher Scientific, CAT#: AAA1390018); (S)-(+)-Carvone (Fisher Scientific, CAT#: AAL0713014);Citral (Fisher Scientific, CAT#: AAA1616918); (+)-β-Citronellol (Sigma-Aldrich, CAT#: C83201-5G); D-(+)-Fenchone (Fisher Scientific, CAT#: AC366291000); L-(-)-Fenchone (Fisher Scientific, CAT#: AC188110500); (+)-Fenchol (Cayman Chemical, CAT#: 23162); Geraniol (Fisher Scientific, CAT#: AC410900250); (-)-(S)-Limonene (Fisher Scientific, CAT#: AAL1324418); (+)-(R)-Limonene (Fisher Scientific, CAT#: AAL04733AE); (+)-Limonene 1,2-epoxide (Sigma-Aldrich, CAT#: 17763-100ML); (+/-)-(D)-Linalool (Fisher Scientific, CAT#: AC125151000); (-)-Perillyl alcohol (Cayman Chemical, CAT#: 21860); (-)-α-Pinene (Cayman Chemical, CAT#: 21576); (-)-β-Pinene (Cayman Chemical, CAT#: 21577); α-Pinene oxide (Fisher Scientific, CAT#: P136225ML); Sabinene hydrate (Sigma-Aldrich, CAT#: 96573-500MG-F); α-Terpineol (Fisher Scientific, CAT#: AC301610250); Myrtenol (Sigma-Aldrich, CAT#: W343900-100G-K); (S)-cis-Verbenol (Sigma-Aldrich, CAT#: 247065-5G).

### Biosensor Library Generation

Three biosensors were screened in the present work. The DSM library was previously described by Beltrán [1]. Briefly, a double site saturation mutagenesis library comprised saturation mutagenesis (except Cys and Pro) at fifteen positions (59, 83, 89, 92, 94, 108, 117, 120, 122, 141, 159, 160, 163, 164 and 167), and our positions (62, 81, 87 and 110) were restricted to smaller amino acid subsets.

The Terp-shuf. library was made using plasmids with PYR1 terpene-responsive variants from screening the DSM library ($PYR1^{Ger3}$, $PYR1^{Ger4}$, and $PYR1^{Cit1}$) that were combined with the original PYR1 DSM library in a mole ratio of 50/50 (16.7% of each sensor, 50% DSM library), followed by recombinant-based mutagenesis, using nucleotide excision and exchange technology [1]. Resulting shuffled fragments were cloned into the Y2H pBD plasmid by restriction digest followed by ligation.

The third library (HMH) was computationally designed. Since all monoterpenes screened were marked by one major hydrogen bond donor and a large C10 hydrophobic carbon skeleton, we chose to make a deeply mutagenized, hydrophobic PYR1 binding pocket library (HMH). We chose benzylfentanyl as a ligand owing to its hydrophobicity, relatively large size, and a single h bond acceptor. To generate ligand conformers for docking, the 3D conformer of benzylfentanyl was downloaded from PubChem and used as the input file for the confab conformer generation method in the software OpenBabel [2], running all default settings. Ligand conformers were aligned in the PYR1 pocket (PDB ID: 3QN1) along a hydrogen bond acceptor using custom PyRosetta code [3] and visualized in PyMOL [4]. PYR1 sequence mutations likely to contribute to ligand binding were identified by computational design of the protein-ligand complex using Rosetta FastDesign[2] using the default energy function performed using PyRosetta4 version 2021.26+release.b308454c455dd04f6824cc8b23e54bbb9be2cdd7, followed by small to large aliphatic substitutions by visual inspection. Combinations of mutations were encoded in oligo pools (IDT) divided into 3 cassettes of 200bp, 142bp, and 167 bp in length to be cloned into the PYR1 yeast two-hybrid destination vector pJS646 by Golden Gate assembly [1,5,6]. Constructs were transformed by electroporation into *E. coli* XL1-blue and plated on LB-chloramphenicol in large bioassay dishes, along with a serial dilution plate to evaluate transformation efficiency. Transformation plates were then scraped to collect colonies and midiprepped (ZymoPURE II Plasmid Midiprep Kit) to extract plasmid DNA. The final HMH library was pooled as equal parts from two different Golden Gate assemblies and transformations, to increase the overall number of transformants, and library composition was confirmed by Quintara AmpExpress next generation sequencing. All plasmid libraries were transformed into MAV99.

### *S. cerevisiae* Library Cloning

A single colony of MaV99 (pAD-HAB1) was picked and grown in 2 mL SD-Leu with 1 M sorbitol overnight at 30 °C. Next, 1 mL of this starter culture was used to inoculate 50 mL of SD-Leu with 1 M sorbitol in a 50 mL conical tube. After growing to an OD600 of 1.0-1.2 (14-16 hrs at 30 °C), the cells were harvested by centrifugation at 2,000 g for 5 minutes. Cells were resuspended in 20 mL 5X TE, LiOAc (0.5 M, pH 7.5) and shaken at 200 RPM, 30 °C for 45 minutes. Then, 500 uL of dithiothreitol (DTT) was added to the solution and incubated for an additional 15 minutes. Next, 30 mL ice-cold sterile $H_2O$ was added and the solution was centrifuged at 2,000 g, 4 ℃ for 5 minutes. The supernatant was removed and the cell pellet was resuspended in 50 mL cold $H_2O$, the mixture was then centrifuged at 2,000 g, 4 ℃ for 5 minutes. The supernatant was again removed and the pellet was resuspended in 1 mL cold $H_2O$. This was then transferred to a pre-chilled 1.5 mL tube and centrifuged at 6,000 g, 4 ℃ for 1 minute. The supernatant was removed and 1 mL ice-cold 1 M sorbitol was added, mixed gently, and centrifuged at 6,000 g for 1 minute at 4 ℃. The supernatant was again removed and the cell pellet was resuspended in 400 µL ice-cold 1 M sorbitol. Next, 7.5 µg of library plasmid DNA was added and mixed well. This mixture was then incubated on ice for 15 minutes. The mixture was then transferred to a 0.4 cm ice-cold electroporation cuvette and pulsed at 2.5 kV, 25 µF, 200 Ω. Immediately, 1 mL ice-cold 1 M sorbitol was added and the mixture was incubated for 30 minutes at room temperature. The mixture was plated on SD-Trp,-Leu containing 1 M sorbitol. After 3 days of incubation the cells were washed off with 1X TE and plated on SD-Trp,-Leu,+1 g/L 5-fluoroorotic acid and incubated for 3 days. The cells were then washed off again and replated SD-Trp,-Leu,+1 g/L 5-fluoroorotic acid and incubated for an additional 3 days. This step is done to screen out constitutive PYR1 mutants. After incubation the cells were washed off, again with 1X TE, and stored as 30% glycerol stocks until use. A minimum transformation efficiency of 1 million CFU was achieved, typically by pooling two transformations.

### *S. cerevisiae* Transcriptional Circuits

Our biosensor reporting system is based on the Agilent GAL4 Two-Hybrid Phagemid Vector Kits. In short an integrated GAL4-UAS is used to drive conditional reporting of *lacZ* (for β-galactosidase staining) and *URA3* (for screening and growth assays). A two plasmid system is used to drive activation of the GAL4-UAS. The pAD plasmid is maintained by *AmpR* and *LEU2* and contains pADH1-SV40NLS-GAL4AD-HAB1-ADH1t for activating transcription when PYR1 is bound. The pBD plasmid is maintained by *ChlR* and *TRP1* and contains pADH1-SV40NLS-GAL4BD-PYR1-ADH1t for binding to the GAL4-UAS until ligand is present and the HAB1 complex can bind.

### Yeast Two-Hybrid β-Galactosidase Staining and Growth Validation

To determine the limit of detection for each biosensor, a growth assay and β-galactosidase assay were performed. For growth assays, the various biosensor-containing yeast cells were grown as spot assays by suspending a colony in 100 µL of 1X TE and dropping 2 µL onto SD-Trp,-Leu,-Ura plates containing various concentrations of the target ligand. After 3 days of growth the plates were imaged. The β-galactosidase assay was conducted in a similar manner, but grown on SD-Trp,-Leu plates containing various concentrations of the target ligand. After 3 days of growth the cells were submerged in chloroform for 15 minutes, the chloroform was removed, and then were allowed to dry for 10 minutes. An X-gal agarose solution, consisting of Z-buffer (8.5 g/L $Na_2HPO_4$, 4.8 g/L $NaH_2PO_4$, 0.75 g/L KCl, 0.12 g/L $MgSO_4$), agarose (6 g/L), BME (0.27%, added after cooled to 50 °C), and X-gal solution (0.69% of 50 g/L solution in N,N-dimethylformamide, added after cooled to 50 °C), was then gently pipetted over the cells and the plates were incubated overnight at 37 °C. The plates were then imaged.

### *K. marxianus* Transcriptional Circuits

To port PYR1 biosensors to *K. marxianus*, we used pNR007 (HIS3) as a backbone to clone in new biosensors. This plasmid contains the PYR1 (TEF1-SV40NLS-Z4DBD-PYR1-ScTDH1) and HAB1 (ScPGK1p-VP64AD-HAB1-PGK1t) cassettes necessary for binding to the integrated cassette (ABZ1::$Z4_4$-HTB1-reporter-CYC1t) when ligand is present where “reporter” is either eGFP (strain Ys2042 and derivatives) or neoR (strain sNR136 and derivatives) for GFP production or G418 resistance, respectively.

For cloning, pNR007 was digested with EcoRI and BamHI to generate the backbone. PYR1 was amplified from plasmids containing biosensors from *S. cerevisiae* (based on pBD) with the primers sw420 and sw422. Gibson assembly (NEB 2X MM) was used to clone new sensors into the pNR007 backbone. Reporter cassettes were integrated into *K. marxianus* with pSW257 and pIW1134 (for eGFP reporter) or pNR012 and pIW1134 (for neoR reporter).

### GFP Reporter Assay

Cells containing eGFP reporter cassette (strain Ys2042) and PYR1/HAB1 biosensor plasmid (based on pNR007) were used to inoculate 2 mL SD-H media cultures containing 2% glucose and pre-cultured; cells were then passaged into wells of a 96-deep-well plate (USA Scientific, Orlando, FL, USA) in 1 mL media ($OD_{600}$ = 0.05). Up to 5 μL of ligand stocks (solvated in DMSO) were added immediately after inoculation, plates sealed with an AeraSeal film (Excel Scientific, Victorville, CA, USA), and grown at 1,000 rpm, and 90 % humidity for 12 h. The cells were harvested by centrifuge at 5000 g for 3 min. After discarding the supernatant, the cells were washed once with 200 uL sterile water and then resuspended in 200 uL sterile water for flow cytometry analysis. BD accuri™ C6 flow cytometer (BD Bioscience) was used for data collection and analysis. A control cell population without fluorescent protein expression was first run to identify basal cell auto-fluorescence before collecting data for the experimental samples. For each sample, 10,000 events were collected within 5 seconds of read time. The forward scatter, side scatter, and eGFP fluorescence were recorded for each event. All experiments were performed in biological triplicate.

### CRISPR gRNA Library Design

The CBS6556 genome sequence (GenBank assembly GCA_016625955.1) was annotated by an ab initio gene predictor AUGUSTUS [7], which was independently trained by gene models of two well-known *K. marxianus* strains, DMKU3-1042 and FIM1. The 10-fold coverage sgRNA library was designed using CHOPCHOP v3 [8] to target the first 5% to 65% of every CDS based on each annotation prediction. The design process was similar to our previously published paper [9]. In brief, custom Python scripts determined the start and end locations for the targeted regions, which were then input into the CHOPCHOP command-line script. CHOPCHOP provided a list of potential sgRNAs for each CDS, including quality and efficiency parameters. We further enhanced the library with a custom Python script that added a uniqueness measure, Seed_MM0, to minimize off-target effects by counting the number of off-targets with 0 mismatches in the seed region (the last 12 bp of the sgRNA near the PAM motif). sgRNAs designed for each CDS were ranked based on their uniqueness and their cumulative targeting efficiency predictions and. The best 10 sgRNAs targeting each CDS were then chosen to be included in the final library. Two separate libraries were designed using annotation predictions obtained from DMKU3-1042 and FIM1. The final library designed for CBS6556 was the union of these two libraries.

### *K. marxianus* Genome-Wide Transformation

Genome-wide transformations were performed in the same way as individual transformations except for some details. To ensure sufficient library coverage, cells were first transformed with the PYR1/HAB1 biosensor plasmid (pNR067). High transformation efficiency was harder to achieve in this strain due to multiple gene integrations and the burden of maintaining a plasmid. Sixteen experimental transformations were performed as well as two controls with an empty vector (pIW1213) into our screening strain. Since the strain we transformed contained Cas9, our apparent transformation efficiency would be lower with the library than with the empty vector due to some guides causing a cut to an essential gene, leading to lower viability or cell death. For our experimental transformations we obtained an average of $1.7x10^4$ transformants per reaction and for our empty vector control we obtained an average of $1.2x10^5$ transformants per reaction.

### Knockout Validation

Due to the gRNA library being on a low copy number plasmid (pIW1213 backbone), yeast plasmid extraction was required to obtain sufficient DNA to amplify the gRNA region. Zymoprep Yeast Plasmid Miniprep II kits were used on all picked hits to extract the gRNA plasmid. POL3_gRNAseq_for and POL3_gRNAseq_rev primers were used to amplify the gRNA region and sanger sequence. If a hit was found to contain a targeting, whole guide (5% of colonies picked), ~600 bp of the guide target site was amplified with colony PCR and sent for sanger sequencing (**Table S3**). Successful gene deletion was confirmed if a frameshift mutation was observed that caused a premature stop codon.

### Terpene Production and GC-FID Validation

Starter cultures were prepared by inoculating 2 mL of YPD with a single colony and growing overnight at 30 °C. A 250 mL flask with 2 mL of dodecane overlay and 50 mL of YPD was inoculated to a starting OD600 of 0.05 and grown at 30 °C for 16 hrs. Ten grams of sodium chloride was added to the flask and allowed to shake for an additional 15 minutes to break any emulsion that may have formed. The entire culture was collected in a 50 mL centrifuge tube and spun down at 10,000 g for 3 minutes. The dodecane layer was carefully collected into a 200 µL insert in a GC vial for GC-FID analysis

GC-FID analysis of terpenes was performed on an Agilent GC-2010 with a 30 m 0.32 mm ID DB-Wax column. Sample was injected at a volume of 5 µL with a splitter temperature of 200 °C and a starting oven temperature of 50 °C. After 3 minutes at 50 °C, the oven temperature ramped at a rate of 5 °C/min to a final temperature of 220 °C and held for 3 minutes. Then, the temperature was ramped back down to 50 °C at a rate of 40 °C/min. Geranic acid eluted at 23.0 min and geraniol at 24.7 min. Calibration curves were made with 0.2, 0.5, 1.0, 2.0, and 5.0 mg/L geraniol and geranic acid in duplicate with an R-squared of >0.98. Concentration of terpenes found in dodecane was divided by 25 to account for the volume of media used. Extraction efficiencies were determined by shaking YPD + dodecane flasks with the aforementioned experimental conditions with 200 µg/L geraniol or geranic acid, but no cells. Extraction efficiency for geraniol was 55% (±10.3%) and for geranic acid was 95% (±5.61%). Dodecane titers were divided by these extraction efficiencies to account for terpenes lost in the aqueous phase. B-citronellol extraction efficiency was 82% (±13.9%), but no detectable quantities were found in any samples.

### Table S1: Strains

| Strain | Description | Source |
|---|---|---|
| E. coli TOP 10 | F- mcrA Δ (mrr-hsdRMS-mcrBC) Φ80lacZΔM15 Δ lacX74 recA1 araD139 Δ (araleu)7697 galU galK rpsL (StrR) endA1 nupG | Thermo Fisher Scientific |
| MaV99 (*Sc*) | MATa trp1-901 leu2-3,112 hisΔ200 ade2-101 gal4Δ gal80Δ can1rcyh2rLYS2::(GAL1::HIS3) GAL1::lacZ SPO13::10xGAL4site::URA3 | M. Vidal et al.[10] |
| Ys626 | *K. marxianus* CBS6556 ΔURA3 ΔKat1 ΔAlpha3 | Löbs et al.[11] |
| Ys2042 | YS626 abz1::Z4BS-HTB1core-GFP-CYC1t | Daffern et al.[12] |
| FS_E | K. marxianus CBS6556 ΔURA3 ΔKat1 ΔAlpha3 I4::PScTEF1-KmCas9-SV40-ScCYC1t | This work |
| sNR116 | XYL2::GPDp-tCrGES-CYCt dURA stable haploid | This work |
| sNR138 | URA3d HIS3d stable haploid (Ys626 HIS3d) | This work |
| sNR139 | XYL2::GPDp-tCrGES-CYCt dURA dHIS stable haploid | This work |
| sNR140 | URA3d::Z4(4)-UAS-KanR-CYCt stable haploid dHIS | This work |
| sNR141 | URA3d::Z4(4)-UAS-KanR-CYCt stable haploid dHIS | This work |
| sNR143 | stable haploid dURA dHIS I4::Cas9 | This work |
| sNR144 | stable haploid dURA dHIS I4::Cas9; XYL2::GPDp-tCrGES-CYCt | This work |
| sNR145 | stable haploid dURA dHIS I4::Cas9; URA3d::Z44-UAS-KanR-CYCt | This work |
| sNR146 | stable haploid dURA dHIS I4::Cas9; URA3d::Z44-UAS-KanR-CYCt; XYL2::GPDp-tCrGES-CYCt | This work |
| sNR184 | URA3d HIS3d stable haploid KLMA_10600 KO | This work |
| sNR185 | URA3d HIS3d stable haploid CDC25 KO | This work |
| sNR186 | URA3d HIS3d stable haploid CTT1 KO | This work |
| sNR187 | URA3d HIS3d stable haploid RRD1 KO | This work |
| sNR188 | URA3d HIS3d stable haploid CLN3 KO | This work |
| sNR189 | URA3d HIS3d stable haploid YSP2 KO | This work |
| sNR190 | URA3d HIS3d stable haploid ALD4 KO | This work |
| sNR191 | URA3d HIS3d stable haploid KLMA_40017 KO | This work |
| sNR192 | URA3d HIS3d stable haploid YAP1 KO | This work |
| sNR193 | URA3d HIS3d stable haploid CWH43 KO | This work |
| sNR194 | URA3d HIS3d stable haploid TIP41 KO | This work |
| sNR195 | XYL2::GPDp-tCrGES-CYCt dURA dHIS stable haploid KLMA_10600 KO | This work |
| sNR196 | XYL2::GPDp-tCrGES-CYCt dURA dHIS stable haploid CDC25 KO | This work |
| sNR197 | XYL2::GPDp-tCrGES-CYCt dURA dHIS stable haploid CTT1 KO | This work |

| | | |
|---|---|---|
| sNR198 | XYL2::GPDp-tCrGES-CYCt dURA dHIS stable haploid ARO4 KO | This work |
| sNR199 | XYL2::GPDp-tCrGES-CYCt dURA dHIS stable haploid CLN3 KO | This work |
| sNR200 | XYL2::GPDp-tCrGES-CYCt dURA dHIS stable haploid YSP2 KO | This work |
| sNR201 | XYL2::GPDp-tCrGES-CYCt dURA dHIS stable haploid ALD4 KO | This work |
| sNR202 | XYL2::GPDp-tCrGES-CYCt dURA dHIS stable haploid KLMA_40017 KO | This work |
| sNR203 | XYL2::GPDp-tCrGES-CYCt dURA dHIS stable haploid YAP1 KO | This work |
| sNR204 | XYL2::GPDp-tCrGES-CYCt dURA dHIS stable haploid CWH43 KO | This work |
| sNR205 | XYL2::GPDp-tCrGES-CYCt dURA dHIS stable haploid TIP41 KO | This work |
| sCLM_T1 | MaV99+pAD+pBD (-)-Perillyl Alcohol 10 | This work |
| sCLM_T2 | MaV99+pAD+pBD (+/-)-Linalool 1 | This work |
| sCLM_T3 | MaV99+pAD+pBD (+/-)-Linalool 2 | This work |
| sCLM_T4 | MaV99+pAD+pBD (+/-)-Linalool 3 | This work |
| sCLM_T5 | MaV99+pAD+pBD (+/-)-Linalool 4 | This work |
| sCLM_T6 | MaV99+pAD+pBD (+/-)-Linalool 5 | This work |
| sCLM_T7 | MaV99+pAD+pBD (+/-)-Linalool 7 | This work |
| sCLM_T8 | MaV99+pAD+pBD (+/-)-Linalool 8 | This work |
| sCLM_T9 | MaV99+pAD+pBD (S)-cis-Verbenol 1 | This work |
| sCLM_T10 | MaV99+pAD+pBD (S)-cis-Verbenol 2 | This work |
| sCLM_T11 | MaV99+pAD+pBD (S)-cis-Verbenol 5 | This work |
| sCLM_T12 | MaV99+pAD+pBD (S)-cis-Verbenol 6 | This work |
| sCLM_T13 | MaV99+pAD+pBD (S)-cis-Verbenol 7 | This work |
| sCLM_T14 | MaV99+pAD+pBD (S)-cis-Verbenol 8 | This work |
| sCLM_T15 | MaV99+pAD+pBD α-Terpineol 1 | This work |
| sCLM_T16 | MaV99+pAD+pBD α-Terpineol 2 | This work |
| sCLM_T17 | MaV99+pAD+pBD α-Terpineol 4 | This work |
| sCLM_T18 | MaV99+pAD+pBD Geraniol 1/(-)-Perillyl Alcohol 10 | This work |
| sCLM_T19 | MaV99+pAD+pBD Geraniol 10 | This work |
| sCLM_T20 | MaV99+pAD+pBD Geraniol 4 | This work |
| sCLM_T21 | MaV99+pAD+pBD Geraniol 11 | This work |
| sCLM_T22 | MaV99+pAD+pBD Geraniol 12 | This work |
| sCLM_T23 | MaV99+pAD+pBD Geraniol 13 | This work |
| sCLM_T24 | MaV99+pAD+pBD Geraniol 14 | This work |
| sCLM_T25 | MaV99+pAD+pBD Geraniol 15 | This work |

| | | |
|---|---|---|
| sCLM_T26 | MaV99+pAD+pBD Geraniol 21 | This work |
| sCLM_T27 | MaV99+pAD+pBD Geraniol 22 | This work |
| sCLM_T28 | MaV99+pAD+pBD Geraniol 23 | This work |
| sCLM_T29 | MaV99+pAD+pBD Myrtenol 1 | This work |
| sCLM_T30 | MaV99+pAD+pBD Myrtenol 2 | This work |
| sCLM_T31 | MaV99+pAD+pBD Myrtenol 3 | This work |
| sCLM_T32 | MaV99+pAD+pBD Myrtenol 6 | This work |
| sCLM_T33 | MaV99+pAD+pBD Myrtenol 7 | This work |
| sCLM_T34 | MaV99+pAD+pBD Myrtenol 8 | This work |
| sCLM_T35 | MaV99+pAD+pBD R-(-)-Carvone 1 | This work |
| sCLM_T36 | MaV99+pAD+pBD R-(-)-Carvone 2 | This work |
| sCLM_T37 | MaV99+pAD+pBD R-(-)-Carvone 3 | This work |
| sCLM_T38 | MaV99+pAD+pBD β-Citronellol 1 | This work |
| sCLM_T39 | MaV99+pAD+pBD β-Citronellol 2 | This work |
| sCLM_T40 | MaV99+pAD+pBD β-Citronellol 3 | This work |
| sCLM_T41 | MaV99+pAD+pBD β-Citronellol 4/Geraniol 20 | This work |
| sCLM_T42 | MaV99+pAD+pBD β-Citronellol 5 | This work |
| sCLM_T43 | MaV99+pAD+pBD β-Citronellol 6 | This work |
| sCLM_T44 | MaV99+pAD+pBD β-Citronellol 8 | This work |
| sCLM_T45 | MaV99+pAD+pBD β-Citronellol 9 | This work |

### Table S2: Plasmids

| Plasmid | Description | Source |
|---|---|---|
| pIW601 | PScTEF1-KmCas9-SV40-ScCYC1t, PKmPRP1-tRNAGly-PspXI recognition site-SUP4 (Km KO empty vector) | Löbs et al.[11] |
| pIW538 | pIW601with URA3 targeting sgRNA | Li et al.[13] |
| pIW447 | pIW601with XYL2 targeting sgRNA | Li et al.[13] |
| pIW1198 | 700bp URA3 up and downstream, pIW578 with EGFP | Li et al.[13] |
| pIW1136 | 700bp XYL2 up and downstream, pIW578 with EGFP | Li et al.[13] |
| SW286 | Reporter plasmid (used for cloning) | Wei et al.[14] |
| SW310 | PYR1/HAB1 Km biosensor plasmid ABA (wt) (URA3) | Wei et al.[14] |
| pACT-HAB1 | pACT with GAL4 AD-HAB1 for Y2H experiments | Park et al.[15] |
| pBD-PYR1 | pBD with GAL4 BD-PYR1 (wt) for Y2H experiments | Park et al.[15] |
| pNR003 | pIW1136; GFP replaced with tCrGES | This work |
| pNR012 | KanR reporter integration plasmid into ABZ1 (used for cloning) | This work |
| pNR031 | pIW1198; GFP replaced with NeoR reporter | This work |
| pNR007 | PYR1/HAB1 Km biosensor plasmid ABA (wt) (HIS3) | This work |
| pNR010 | pNR007 Ger3 sensor | This work |
| pNR065 | pNR007 Ger11 sensor | This work |
| pNR066 | pNR007 Ger14 sensor | This work |
| pNR067 | pNR007 Ger15 sensor | This work |
| pNR068 | pNR007 Ger21 sensor | This work |
| pNR069 | pNR007 Ger22 sensor | This work |
| pNR070 | pIW601 with KLMA_10600 gRNA | This work |
| pNR071 | pIW601 with CDC25 gRNA | This work |
| pNR072 | pIW601 with CTT1 gRNA | This work |
| pNR073 | pIW601 with ARO4 gRNA | This work |
| pNR075 | pIW601 with CLN3 gRNA | This work |
| pNR076 | pIW601 with YSP2 gRNA | This work |
| pNR078 | pIW601 with ALD4 gRNA | This work |
| pNR079 | pIW601 with KLMA_40017 gRNA | This work |
| pNR080 | pIW601 with YAP1 gRNA | This work |
| pNR081 | pIW601 with CWH43 gRNA | This work |
| pNR082 | pIW601 with TIP41 gRNA | This work |

### Table S3: Primers

| Primer | Sequence | Description |
|---|---|---|
| 1198BB_4repfor | GTCATGTAATTAGTTATGTCACGC | 1198 backbone for KanR reporter |
| 1198BB_4reprev | GCATTAACAACCCTCTAGG | 1198 backbone for KanR reporter |
| Rep4_1198for | GCCGAGGAAGTTACGAAAGAACCTAGAGGGTTGTTAATGCcgcaaaactagtctcatgc | Amplify KanR reporter from pNR012 for 1198 |
| Rep4_1198rev | GAGGGCGTGAATGTAAG | Amplify KanR reporter from pNR012 for 1198 |
| 257neoBB_for | GCTCGAGTCATGTAATTAGTTATGTCAC | Linearize sw257 to make pNR012 |
| 257neoBB_rev | GCTTTTTCTGTATTATTTGGTTATCTGGG | Linearize sw257 to make pNR012 |
| neoR257_for | TACTACGCCCCCCAGATAACCAAATAATACAGAAAAAGCCCATGAGCCATATTCAACGGG | Amplify KanR from sw257 |
| neoR257_rev | AAGCGTGACATAACTAATTACATGACTCGAGCGTTAGAAAAACTCATCGAGCATCAAATG | Amplify KanR from sw257 |
| sw918 | AACCTCTGACACATGCAGCTCTAGTACACTCTATATTTTTTTATGCCT | Amplify HIS3 from SW286 into SW310 to make pNR007 |
| sw919 | TTTGTGAGTTTAGTATACATGCATCTACATAAGAACACCTTTGGTGG | Amplify HIS3 from SW286 into SW310 to make pNR007 |
| sw420 | ACACAAAGATACATACGGGAGGATCCATGCCTTCGGAGTTAACACCAG | Amplify PYR1 sensors from pBD into pNR007 |
| sw422 | CCTCATCAAGATTGCTTTATGCTAGCTCACGTCACCTGAGAACCACTTC | Amplify PYR1 sensors from pBD into pNR007 |
| T24-27_for | gcaccttcaccaagtcg | KLMA_10600 colony PCR primer |
| T24-54_for | cacaggggtccatactgg | CDC25 colony PCR primer |
| T24-57_for | ccatctgatccgccagtttattc | CTT1 colony PCR primer |
| T24-70_for | ggttgtttcgaaacttatgcatcc | ARO4 colony PCR primer |
| T24-73_for | cctgaaattacttatcagatgagaagcc | CLN3 colony PCR primer |
| T24-76_for | cgattaagcaagatggcgc | YSP2 colony PCR primer |
| T28-2_for | GCAAAGGGTGATGTTGCTC | ALD4 colony PCR primer |
| T28-26_for | caaaacggttcaaccaatctttg | KLMA_40017 colony PCR primer |
| T28-70_for | gtggatatgccgttgctg | YAP1 colony PCR primer |
| T28-81_for | gctgctagcccatcaaag | CWH43 colony PCR primer |
| T28-90_for | gatccctgcgggacaac | TIP41 colony PCR primer |
| T24-27_rev | cctacagcgcatctcagtg | KLMA_10600 colony PCR primer |

| | | |
|---|---|---|
| T24-54_rev | ccatcaaggacacgaatagc | CDC25 colony PCR primer |
| T24-57_rev | ccttccttgttcacaaacttgaac | CTT1 colony PCR primer |
| T24-70_rev | gacaacactttcaacatcaacaaagg | ARO4 colony PCR primer |
| T24-73_rev | cgctctcattaatactctccaagatag | CLN3 colony PCR primer |
| T24-76_rev | gtaggtgacaccactgtgtc | YSP2 colony PCR primer |
| T28-2_rev | GAACCGGCACAGCAAAC | ALD4 colony PCR primer |
| T28-26_rev | ccgattcatacgtagcctactttg | KLMA_40017 colony PCR primer |
| T28-70_rev | ggaattagcacgatgagcac | YAP1 colony PCR primer |
| T28-81_rev | gcgcagagctaggctttc | CWH43 colony PCR primer |
| T28-90_rev | cgtagtttggcgagcc | TIP41 colony PCR primer |
| URA3d_for | GCATTTCGAAACAAGAGGGG | URA3d colony PCR primer |
| URA3d_rev | GTTCTAGCAATCGACGGTCC | URA3d colony PCR primer |
| XYL2_for | GAAGATGTGCTATACTGTGGATAAGG | XYL2 colony PCR primer |
| XYL2_rev | CATCGATTGCTTCGTCAAACTTG | XYL2 colony PCR primer |
| pBD_Fw2 | CGGAAGAGAGTAGTAACAAAGGT | PYR1 insert PCR primer |
| pBD_Rev2 | GAAATTCGCCCGGAATTAGCTTG | PYR1 insert PCR primer |

### Table S4: Guide RNAs

| Guide Name | Sequence | Gene Target |
|---|---|---|
| T24-27 | GAACTGCGACATGATAGCCG | KLMA_10600 |
| T24-54 | AAAGATAGCATGCATACTGG | CDC25 |
| T24-57 | GCATCGCGAATGAAAAACAC | CTT1 |
| T24-70 | GTGTTCATTACCCTTGGTGG | ARO4 |
| T24-73 | CTCTGCAAGGTGGGTAACGA | CLN3 |
| T24-76 | TCTTGGTCTGACATCAACCG | YSP2 |
| T28-2 | GAAACGTACAATGCAGACAA | ALD4 |
| T28-26 | ACTATAAAACAAGAACACGA | KLMA_40017 |
| T28-70 | GGAAAGCCCTCTGTGCCGCA | YAP1 |
| T28-81 | GTAAGGACTGTTACATGTGG | CWH43 |
| T28-90 | GATGATATTTGGTAATTCTG | TIP41 |
| URA3d | TTGCCGAGTTATCGTCCAAG | URA3d |
| XYL2 | TGGTCAAGGTTGGCGATCGT | XYL2 |

**Table S5.** HMH library design. The library was designed as individual cassettes covering linear stretches of the PYR1 gene. Library assembly was performed using Golden Gate.

| Cassette | Position | WT AA | Library |
|---|---|---|---|
| 1 | 59 | K | KRQ |
| | 83 | V | VW |
| | 87 | L | LAV |
| | 89 | A | AG |
| | 92 | S | SFIM |
| | 94 | E | EKQ |
| 2 | 108 | F | FAV |
| | 117 | L | LAV |
| | 120 | Y | YAFV |
| | 122 | S | SGL |
| 3 | 141 | E | EK |
| | 159 | F | FAILMT |
| | 160 | A | AIMV |
| | 163 | V | AFWMY |
| | 167 | N | NA |

**Table S6.** Theoretical coverage of HMH library.

| | |
|---|---|
| Theoretical library size | 438,944 |
| total # *E. coli* transformants | 4.0e6 |
| Estimated library coverage | >99% |

β-Galactosidase Staining

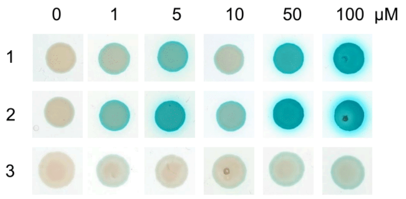


PYR1$^{R\text{-}(-)\text{-}Carvone}$

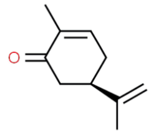


Sequences

| | 59 | 62 | 83 | 89 | 92 | 108 | 117 | 122 | 160 | 163 | LOD (μM) | Library |
|---|---|---|---|---|---|---|---|---|---|---|---|---|
| (R)-(-)-Carvone1 | **Q** | **V** | **W** | **G** | S | **V** | **V** | S | A | **F** | 1 | HMH |
| (R)-(-)-Carvone2 | **Q** | I | **W** | **G** | S | **V** | **V** | S | A | **F** | 1 | HMH |
| (R)-(-)-Carvone3 | **Q** | I | V | **G** | **M** | **V** | L | **L** | **V** | **F** | 1 | HMH |

β-Galactosidase Staining

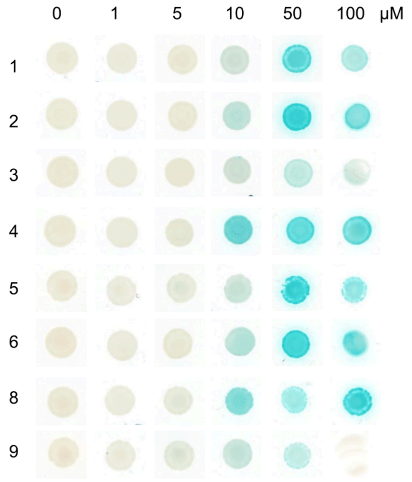


PYR1$^{\beta\text{-}Citronellol}$

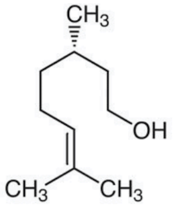


Sequences

| | 81 | 83 | 87 | 120 | 122 | 159 | 160 | 167 | LOD (μM) | Library |
|---|---|---|---|---|---|---|---|---|---|---|
| β-Citronellol1 | V | V | L | **V** | S | F | A | N | 10 | DSM |
| β-Citronellol2 | V | V | L | **V** | S | **V** | A | N | 10 | Terp-shuf. |
| β-Citronellol3 | V | **I** | L | Y | S | F | A | **T** | 10 | Terp-shuf. |
| β-Citronellol4 | V | V | L | Y | S | **L** | **M** | N | 10 | Terp-shuf. |
| β-Citronellol5 | V | V | L | Y | S | **V** | A | **E** | 10 | Terp-shuf. |
| β-Citronellol6 | V | V | **V** | Y | S | **L** | A | N | 10 | Terp-shuf. |
| β-Citronellol8 | V | V | L | Y | **G** | **I** | A | N | 10 | Terp-shuf. |
| β-Citronellol9 | **I** | V | L | **H** | S | F | A | N | 10 | Terp-shuf. |

### β-Galactosidase Staining

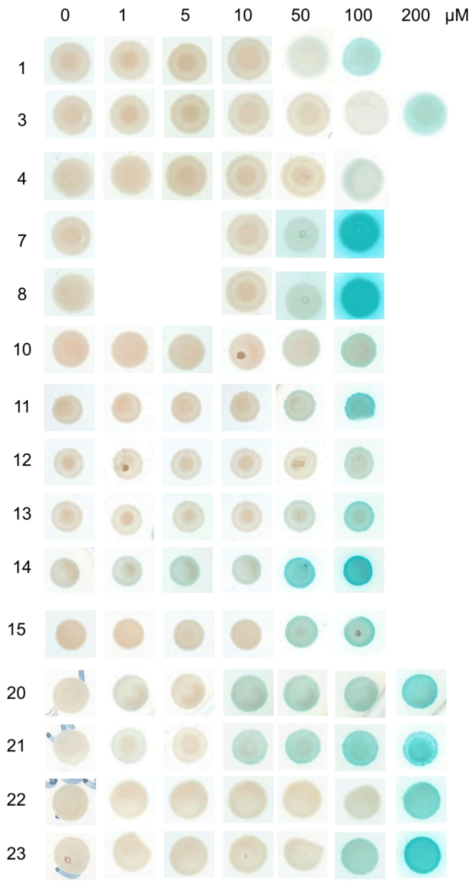


PYR1$^{Geraniol}$

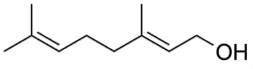


#### Sequences

| | 59 | 81 | 83 | 87 | 89 | 92 | 108 | 120 | 122 | 159 | 160 | 163 | 167 | LOD (μM) | Library |
|---|---|---|---|---|---|---|---|---|---|---|---|---|---|---|---|
| Geraniol1 | K | V | V | L | A | S | F | Y | S | **I** | A | V | N | 50 | DSM |
| Geraniol3 | K | V | V | L | A | S | F | **S** | S | **I** | A | V | N | 200 | DSM |
| Geraniol4 | K | V | V | L | A | S | F | **G** | S | **V** | A | V | N | 100 | DSM |
| Geraniol7 | K | **L** | V | L | A | S | F | Y | S | **I** | A | V | N | 50 | Terp-shuf. |
| Geraniol8 | K | V | V | L | A | S | F | Y | S | **I** | A | **I** | N | 50 | Terp-shuf. |
| Geraniol10 | K | V | V | L | A | S | F | Y | S | **I** | A | **V** | N | 50 | DSM |
| Geraniol11 | **Q** | V | V | **V** | A | S | F | **A** | **L** | **L** | A | V | **A** | 50 | HMH |
| Geraniol12 | K | V | **A** | L | A | S | F | **A** | S | **I** | A | V | N | 100 | HMH |
| Geraniol13 | K | **I** | V | L | A | S | **V** | Y | **G** | **L** | A | V | N | 50 | HMH |
| Geraniol14 | K | V | V | **V** | A | **F** | **A** | Y | **G** | **M** | **V** | V | N | 1 | HMH |
| Geraniol15 | **Q** | V | V | **V** | A | **F** | F | **F** | S | **M** | **M** | V | N | 50 | HMH |
| Geraniol20 (β-Citronellol 4) | K | V | V | L | A | S | F | Y | S | **L** | **M** | V | N | 1 | DSM |
| Geraniol21 | K | V | V | L | A | S | F | **T** | S | **I** | A | V | N | 1 | DSM |
| Geraniol22 | **Q** | V | V | **V** | A | S | **V** | **A** | S | **L** | A | V | N | 200 | HMH |
| Geraniol23 | **Q** | V | **W** | L | **G** | S | F | **A** | S | **L** | A | V | N | 100 | HMH |

### PYR1$^{(+/-)\text{-Linalool}}$

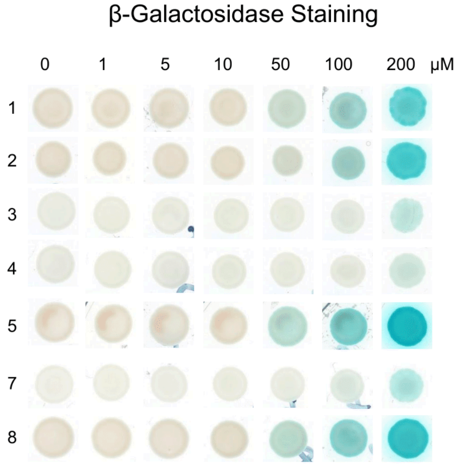


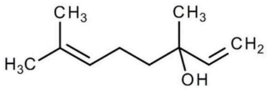


#### Sequences

| | 59 | 62 | 87 | 89 | 92 | 108 | 120 | 122 | 159 | 160 | 167 | LOD (μM) | Library |
|---|---|---|---|---|---|---|---|---|---|---|---|---|---|
| (+/-)-Linalool1 | **Q** | I | **V** | A | S | **V** | Y | **L** | **L** | **L** | N | 50 | HMH |
| (+/-)-Linalool2 | **R** | I | **V** | A | S | F | Y | S | **L** | **M** | N | 100 | HMH |
| (+/-)-Linalool3 | **Q** | I | **V** | **G** | **F** | F | **F** | S | F | **M** | N | 200 | HMH |
| (+/-)-Linalool4 | **Q** | **V** | **V** | **G** | **I** | F | Y | S | **L** | **V** | N | 200 | HMH |
| (+/-)-Linalool5 | **Q** | I | **A** | A | **F** | F | **F** | S | **L** | **L** | **A** | 50 | HMH |
| (+/-)-Linalool7 | **Q** | I | **V** | **G** | **I** | F | **F** | S | **M** | A | N | 200 | HMH |
| (+/-)-Linalool8 | **Q** | I | **A** | A | **F** | F | **F** | S | **M** | **I** | N | 50 | HMH |

### PYR1$^{\text{Myrtenol}}$

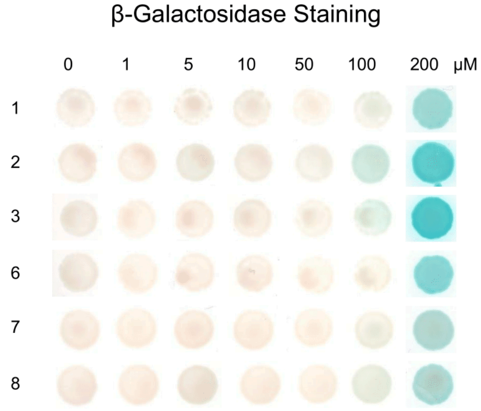


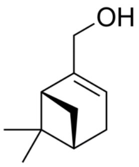


#### Sequences

| | 59 | 81 | 87 | 89 | 92 | 108 | 110 | 120 | 122 | 141 | 159 | 160 | 167 | LOD (μM) | Library |
|---|---|---|---|---|---|---|---|---|---|---|---|---|---|---|---|
| Myrtenol1 | **Q** | V | **V** | **G** | **I** | **V** | I | **A** | S | E | **L** | **M** | N | 200 | HMH |
| Myrtenol2 | **Q** | V | L | **G** | **I** | F | I | **A** | S | E | **L** | A | N | 100 | HMH |
| Myrtenol3 | **Q** | V | **V** | **G** | S | F | I | **A** | S | E | **M** | A | N | 100 | HMH |
| Myrtenol6 | **Q** | **I** | L | A | S | **V** | I | **A** | S | E | F | **V** | N | 200 | HMH |
| Myrtenol7 | **Q** | V | L | A | S | **I** | **L** | Y | **L** | **K** | F | **F** | **A** | 200 | HMH |
| Myrtenol8 | **Q** | V | **V** | **G** | **I** | **V** | I | **A** | S | **K** | **L** | **I** | N | 200 | HMH |

### PYR1$^{(-)\text{-Perillyl Alcohol}}$

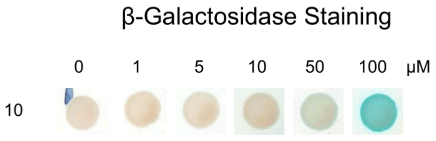


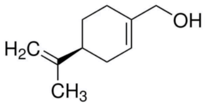


#### Sequences

| | 120 | 159 | LOD (μM) | Library |
|---|---|---|---|---|
| (−)-Perillyl Alcohol10 | **A** | **L** | 100 | HMH |

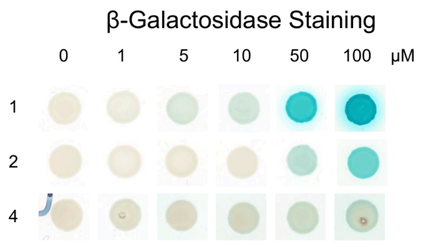


PYR1$^{α\text{-Terpineol}}$

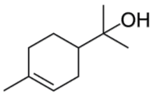


Sequences

| | 59 | 63 | 83 | 87 | 89 | 92 | 108 | 120 | 122 | 159 | 160 | 163 | 164 | LOD (μM) | Library |
|---|---|---|---|---|---|---|---|---|---|---|---|---|---|---|---|
| α-Terpineol1 | K | K | V | **F** | A | S | F | Y | S | F | A | V | **H** | 5 | Terp-shuf. |
| α-Terpineol2 | **Q** | K | V | **A** | **G** | **M** | **A** | Y | **G** | **M** | **I** | **A** | V | 50 | HMH |
| α-Terpineol4 | **Q** | **N** | **W** | **V** | **G** | S | F | **F** | S | F | **L** | V | V | 50 | HMH |

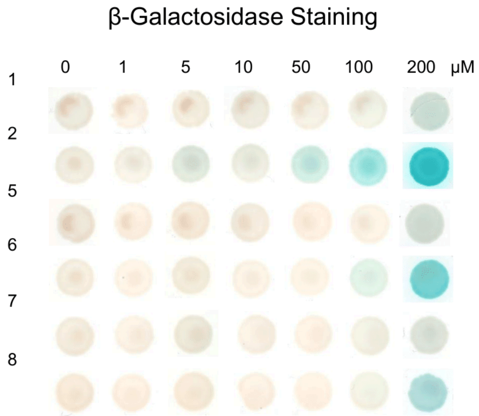


PYR1$^{(S)\text{-cis-Verbenol}}$

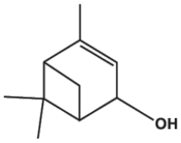


Sequences

| | 59 | 87 | 89 | 92 | 108 | 110 | 117 | 120 | 122 | 141 | 159 | 160 | 163 | 167 | LOD (μM) | Library |
|---|---|---|---|---|---|---|---|---|---|---|---|---|---|---|---|---|
| (S)-cis-Verbenol1 | K | **A** | A | S | F | I | L | Y | S | E | **M** | **M** | V | N | 200 | HMH |
| (S)-cis-Verbenol2 | **Q** | **V** | **G** | **I** | **I** | **V** | L | Y | S | **K** | **L** | **V** | V | N | 5 | HMH |
| (S)-cis-Verbenol5 | K | L | A | S | F | I | **A** | **F** | S | E | F | **L** | V | N | 200 | HMH |
| (S)-cis-Verbenol6 | K | **A** | A | S | F | I | **V** | Y | S | E | F | A | **F** | **A** | 100 | HMH |
| (S)-cis-Verbenol7 | **Q** | **A** | **G** | **I** | **V** | I | L | Y | S | **K** | **L** | **V** | V | N | 200 | HMH |
| (S)-cis-Verbenol8 | **Q** | **V** | **G** | **I** | **V** | I | L | Y | **G** | E | **L** | **I** | V | N | 200 | HMH |

**Figure S1: All Monoterpenoid Biosensors.** Structure of terpenes with blue/white X-gal staining of *S. cerevisiae* spots containing the respective biosensor and reporter circuit (MaV99 Y2H strain). Strains containing the chosen biosensor were spotted as 2 μL drops on SD -trp -leu +ligand plates and allowed to grow for three days at 30 °C, washed with chloroform, and covered in X-gal containing agarose. After one day at 37 °C, staining was obvious and images were taken. Blue indicates activation of the biosensor by the displayed terpene. Amino acid substitutions for all biosensors is also shown. Blue letters indicate an amino acid substitution. Black numbers in the top row indicate the amino acid position in the PYR1 protein sequence.

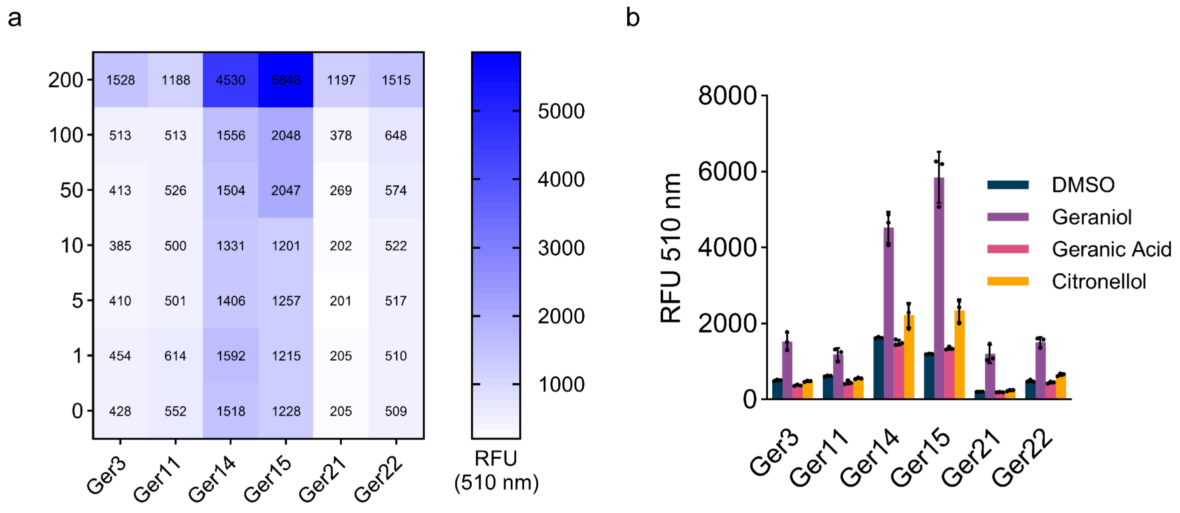


**Figure S2: All geraniol biosensors ported into *K. marxianus*.** GFP fluorescence data for activation of geraniol PYR1 biosensors in *K. marxianus.* The PYR1/HAB1 cassette is carried on a plasmid and the GFP reporter is integrated at *ABZ1*. Overnight cultures were used to inoculate 1 mL wells of SD-HIS liquid culture to an OD600 of 0.05 in a 96-well plate. After 12 hours at 30 °C and 1,000 RPM shaking, fluorescence was measured at 510 nm with flow cytometry. (a) Sensitivity to geraniol at 1, 5, 10, 50, 100, and 200 μM with DMSO as control. (b) Cross reactivity of similar terpenoids at 200 μM with DMSO as control. $PYR1^{Ger21}$ was chosen to be $PYR1^{Geraniol}$.

| | |
|---|---|
| KLMA_10600: | GAACTGCGACATGATAGCCG |
| KO: | GAACTGCGACATGAT - - CCG |
| CDC25: | AAAGATAGCATGCA TACTGG |
| KO: | AAAGATAGCATGCA - - TACTACTGG |
| CTT1: | GCATCGCGAATGAAAAACAC |
| KO: | GCATCGCGAATG - AAAACAC |
| ARO4: | GTGTTCATTACCCTTGG TGG |
| KO: | GTGTTCATTACCCTTGGCGGCATGTCCGGGGTGG |
| CLN3: | CTCTGCAAGGTGGGTAACGA |
| KO: | CTCTGCAAGGTGGGTA - CGA |
| YSP2: | TCTTGGTCTGACATCAACCG |
| KO: | TCTTGGTCTGACATC - ACCG |
| ALD4: | GAAACGTACAATGCAGACAA |
| KO: | GAAACGTACAATGCAGTAACCATAACCCTTTTCCCAACTGTCTCCTAACCTGCTACCTTAATTCCAATACAA |
| KLMA_40017: | ACTATAAAACAAGAACACGA |
| KO: | ACTATAAAACAAGAAC - - GA |
| YAP1: | GGAAAGCCCTCTGTGCC GCA |
| KO: | GGAAAGCCCTCTGTGCCCGCA |
| CWH43: | GTAAGGACTGTTACATGTGG |
| KO: | GTAAGGACTGTTACATG - GG |
| TIP41: | GATGATATTTGGTAA TTCTG |
| KO: | GATGATATTTGGTAATTTCTG |

**Figure S3: Gene knockouts identified in the biosensor-driven genome-wide screen.** For each of the screen hits, the 20 bp sgRNA is displayed and represents the wild-type targeting sequence for each gene. Below each wild-type sequence is the mutation/indel that caused a frame shift premature stop codon downstream of the cut site.

(13) Li, M.; Lang, X.; Moran Cabrera, M.; De Keyser, S.; Sun, X.; Da Silva, N.; Wheeldon, I. CRISPR-Mediated Multigene Integration Enables Shikimate Pathway Refactoring for Enhanced 2-Phenylethanol Biosynthesis in Kluyveromyces

Marxianus. *Biotechnol. Biofuels* **2021**, *14* (1), 3.
(14) Wei, S.; Li, M.; Lang, X.; Robertson, N. R.; Park, S.-Y.; Cutler, S. R.; Wheeldon, I. Repurposing Plant Hormone Receptors as Chemically-Inducible Genetic Switches for Dynamic Regulation in Yeast. *Metab. Eng.* **2024**, *83*, 102–109.
(15) Park, S.-Y.; Fung, P.; Nishimura, N.; Jensen, D. R.; Fujii, H.; Zhao, Y.; Lumba, S.; Santiago, J.; Rodrigues, A.; Chow, T.-F. F.; Alfred, S. E.; Bonetta, D.; Finkelstein, R.; Provart, N. J.; Desveaux, D.; Rodriguez, P. L.; McCourt, P.; Zhu, J.-K.; Schroeder, J. I.; Volkman, B. F.; Cutler, S. R. Abscisic Acid Inhibits Type 2C Protein Phosphatases via the PYR/PYL Family of START Proteins. *Science* **2009**, *324* (5930), 1068–1071.
